# CMTM4 is an adhesion modulator that regulates skeletal patterning and primary mesenchyme cell migration in sea urchin embryos

**DOI:** 10.1101/2024.10.03.616569

**Authors:** Abigail E. Descoteaux, Marko Radulovic, Dona Alburi, Cynthia A. Bradham

## Abstract

MARVEL proteins, including those of the CMTM gene family, are multi-pass transmembrane proteins that play important roles in vesicular trafficking and cell migration; however, little is understood about their role in development, and their role in skeletal patterning is unexplored. CMTM4 is the only CMTM family member found in the developmental transcriptome of the sea urchin *Lytechinus variegatus*. Here, we validate that LvCMTM4 is a transmembrane protein and show that perturbation of CMTM4 expression via zygotic morpholino or mRNA injection perturbs skeletal patterning, resulting in loss of secondary skeletal elements and rotational defects. We also demonstrate that normal levels of CMTM4 are required for normal PMC migration and filopodial organization, and that these effects are not due to gross mis-specification of the ectoderm. Finally, we show that CMTM4 is sufficient to mediate PMC cell-cell adhesion. Taken together, these data suggest that CMTM4 controls PMC migration and biomineralization via adhesive regulation during sea urchin skeletogenesis. This is the first discovery of a functionally required adhesive gene in this skeletal patterning system.

**Graphical Abstract:** 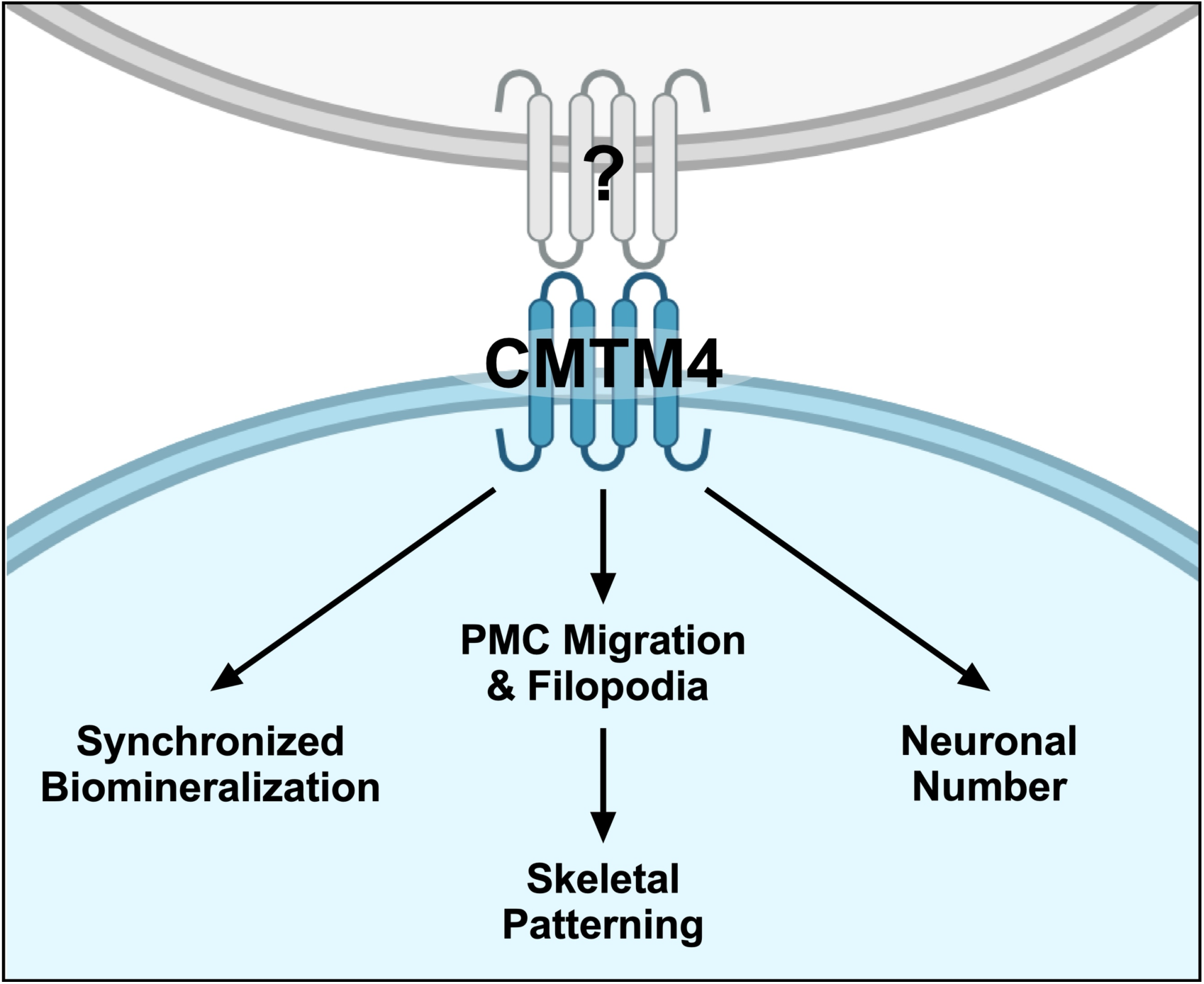

## Introduction

CMTM4 is part of the CMTM gene superfamily, which consists of chemokine-like factor (CKLF) and several CMTM genes: multi-pass transmembrane proteins which contain MARVEL (**m**yelin and lymphocyte **a**nd **r**elated proteins for **ve**sicle trafficking and membrane **l**ink) domains (Sánchez-Pulido et al., 2002; Han et al., 2003; Plate et al., 2010). MARVEL domains are characterized by M-shaped transmembrane domains comprised of four alpha-helices. Proteins containing these domains have been linked to a variety of membrane-associated functions. For example, several MARVEL proteins have been identified at tight junctions (Furuse et al., 1993; Ikenouchi et al., 2005; Raleigh et al., 2010). In mammals, numerous MARVEL domain-containing proteins have also been identified within the membranes of intracellular transport vesicles, the apical plasma membrane, and the endoplasmic reticulum (Zacchetti et al., 1995; Haass et al., 1996; Puertollano and Alonso, 1999), suggesting a role in vesicle trafficking as well as exo- and endocytosis. Indeed, several CMTM family members regulate cancer cell growth and migration by facilitating endocytic recycling of cell surface proteins such as epidermal growth factor receptor (EGFR) (Jin et al., 2005; Li et al., 2014; Yuan et al., 2017; Yuan et al., 2020).

CMTM4 is the only CMTM family member found in the transcriptome of the sea urchin *Lytechinus variegatus*. CMTM4 is relatively understudied compared to other CMTM family members. CMTM4 regulates the trafficking of immune-related proteins such as PD-L1, IL-17, and CXCR4 to the plasma membrane and the recycling of VE-cadherin to endothelial adherens junctions (Mezzadra et al., 2017; Chrifi et al., 2019; Takeuchi et al., 2020; Li et al., 2021; Bona et al., 2022; Knizkova et al., 2022). CMTM4 also facilitates cell migration by modulating cell surface localization of receptors for migratory cues and by regulating the cohesiveness of migratory cell clusters (Chrifi et al., 2019; Xue et al., 2019; Bona et al., 2022). However, the role of CMTM4 in regulating developmental patterning has not yet been directly explored.

In this study, we use the sea urchin *L. variegatus* to explore the role of CMTM4 in development, and found that it is necessary for normal patterning. Skeletal patterning of the sea urchin larval skeleton involves a relatively simple two-component system. The calcium carbonate biomineral is secreted by a population of cells called the primary mesenchyme cells (PMCs) that are patterned by cues from the overlying ectoderm that they presumably receive via the dynamic filopodia which they extend throughout their migration (von Ubisch, 1937; Ettensohn, 1990; Malinda and Ettensohn, 1994; Malinda et al., 1995; Miller et al., 1995; Bradham and McClay, 2006; Piacentino et al., 2016a). To generate the skeletal biomineral, the PMCs take up calcium from the surrounding sea water via endocytosis and secrete it as calcium carbonate into the growing spicule within the lumen of their shared syncytial cable (Hodor and Ettensohn, 1998; Wilt et al., 2008; Stumpp et al., 2012; Schatzberg et al., 2015; Vidavsky et al., 2015; Hu et al., 2020). This process initiates during early gastrulation, when the PMCs first migrate into a ring-and-cords pattern and secrete the first biomineral crystals within the PMC clusters (Hodor and Ettensohn, 1998; Peterson and McClay, 2003; Wu et al., 2007; Wilt et al., 2008; Descoteaux et al., 2023; Zuch and Bradham, 2019a). From this primary pattern, the PMCs from the clusters then migrate outward to pattern and secrete the secondary skeleton. The stereotypic four-armed pluteus larval skeleton is achieved by 48 hours post fertilization (hpf) in *L. variegatus* at room temperature (Wolpert and Gustafson, 1961; Gustafson and Wolpert, 1963; Gustafson and Wolpert, 1967; Ettensohn, 2017; Descoteaux et al., 2023).

Coordinated migration of the PMCs and vesicular trafficking of the skeletal biomineral are therefore key processes in sea urchin skeletal patterning. Because MARVEL proteins are known to play a role in similar processes, we were curious whether CMTM4 was involved in these or other developmental processes during sea urchin skeletal patterning and development. Here, we functionally test the effects of LvCMTM4 perturbation on skeletal patterning, PMC migration, and dorsal-ventral ectodermal specification. We use polychrome labeling and a novel adhesion assay to investigate the functional role of CMTM4. We find that CMTM4 perturbation disrupts the normal spatiotemporal dynamics and patterning of the sea urchin larval skeleton and that it is sufficient for PMC cell-cell adhesion.

## Results

From our developmental transcriptome, we found that expression of *Lv-cmtm4* is low for most of embryonic development, with a large increase in expression between late gastrula and early pluteus stages that remains high during late pluteus stage (Fig. 1A). The increase in *cmtm4* expression after late gastrula stage corresponds with both the migration of the PMCs out of the ring-and-cords to produce the secondary skeletal elements and the differentiation of the major neural cells (Wolpert and Gustafson, 1961; Gustafson and Wolpert, 1963; Gustafson and Wolpert, 1967; Peterson and McClay, 2003; Descoteaux et al., 2023; Bradham et al., 2009); thus, CMTM4 expression is temporally well-suited to play a role in sea urchin skeletal patterning and/or neural development. Next, to assess the spatial expression of *Lv-cmtm4* in the larvae, we used hybridization chain reaction fluorescent *in situ* hybridization (HCR FISH) for *cmtm4* in pluteus-stage embryos, when *cmtm4* expression is highest. We found that *cmtm4* is expressed throughout the embryo, with enrichment in the gut and ciliary band at the dorsal-ventral boundary (Fig. 1B).

**Figure 1.**
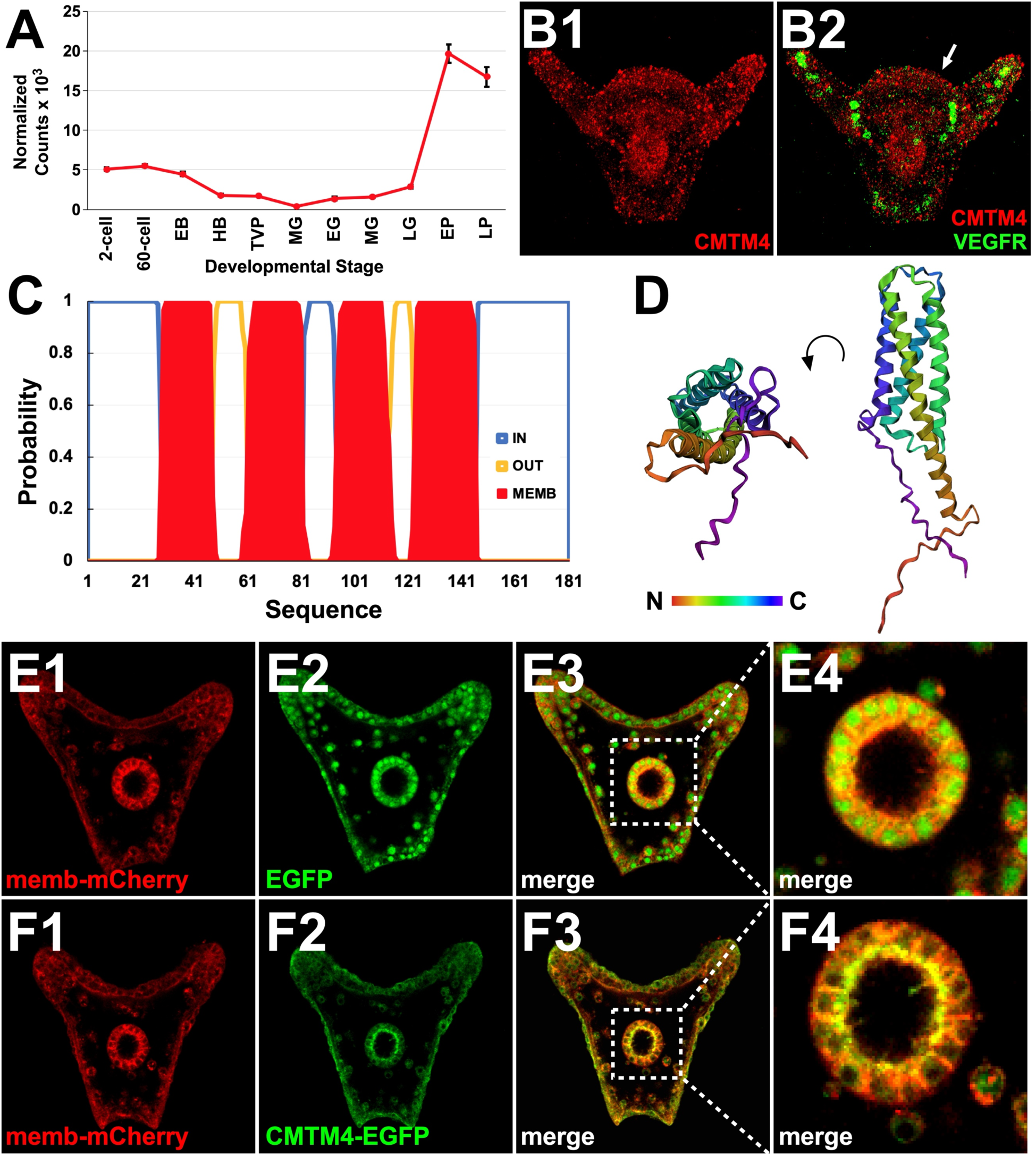
LvCMTM4 is a multi-pass transmembrane protein that is most highly expressed in the gut and ventral ectoderm after late gastrula stage. **A.** Gene expression levels of LvCMTM4 are shown at the indicated developmental stages as normalized read counts ± S.E.M. from temporal transcriptomics data (Hogan et al. 2020). Larval stages are: early blastula (EB, 4 hpf); hatched blastula (HB, 7 hpf); thickened vegetal plate (TVP, 10 hpf); mesenchyme blastula (MB, 13 hpf); early gastrula (EG, 14 hpf); mid-gastrula (MG, 16 hpf); late gastrula (LG, 18 hpf); early pluteus (EP, 36 hpf); and late pluteus (LP, 48 hpf). **B.** HCR FISH exemplars for LvCMTM4 (red) alone (B1) and with VEGFR (green) to mark PMCs (B2) are shown in a 30 hpf control embryo. Arrow in B2 indicates enrichment of LvCMTM4 in the ciliary band. **C.** The intracellular (blue), extracellular (yellow), and transmembrane (red) domains of LvCMTM4 are shown as predicted by DeepTMHMM. **D.** The predicted 3-D protein structure of LvCMTM4 is shown, colorized from N- to C-terminus with the indicated LUT, as predicted by AlphaFold2 in orthogonal views. **E-F.** Confocal slices from live 30 hpf control embryos co-injected with membrane-localized mCherry (1, red) and either EGFP (E2, green) or CMTM4-EGFP (F2, green) are shown individually and merged (3), with the indicated regions (3) magnified (4).

In humans and other species, CMTM4 is a MARVEL domain-containing protein, meaning its protein structure is characterized by four transmembrane domains. We aligned human and *L. variegatus* CMTM4 peptide sequences using UniProt and found that both sequences contain highly hydrophobic internal regions, as would be expected for small, multi-pass transmembrane proteins (Fig. S1) (The UniProt Consortium et al., 2023). To confirm that LvCMTM4 is indeed a MARVEL protein, we used DeepTMHMM to analyze the amino acid sequence of LvCMTM4 and identify possible transmembrane domains (Hallgren et al., 2022). We found that LvCMTM4 is predicted to contain the four transmembrane domains characteristic of MARVEL proteins (Fig. 1C, red). Interestingly, this algorithm also predicted that the N- and C-termini are both intracellular rather than extracellular (Fig. 1C, blue). To further characterize LvCMTM4 protein, we used AlphaFold2 to predict its tertiary structure (Jumper et al., 2021). With this tool, we identified four alpha-helical structures that correspond with the transmembrane domains predicted by DeepTMHMM (Fig. 1D). These alpha-helices appear to be arranged in a pore-like pattern (Fig. 1D).

To assess the cellular localization of LvCMTM4 *in vivo*, we designed a LvCMTM4-EGFP fusion construct in which the stop codon of LvCMTM4 was removed and the amino acid sequence of enhanced green fluorescent protein (EGFP) was added in frame. We co-injected sea urchin zygotes with mRNA generated from this construct along with a membrane-targeted mRNA encoding mCherry. Separately, we co-injected EGFP mRNA with membrane-mCherry mRNA as a negative control. We found that, as expected, EGFP and membrane-mCherry co-injected embryos show red labeling of the membrane and green labeling of the cytosol (Fig. 1E). When LvCMTM4-EGFP mRNA was co-injected with membrane-mCherry mRNA, the green fluorescent signal was instead detected along the cell membranes (Fig. 1F). Since the injection of LvCMTM4-EGFP mRNA at the zygote stage results in global expression of LvCMTM4-EGFP, we expected to see equal CMTM4-EGFP signal throughout the cells; however, interestingly, we find that the CMTM4-EGFP signal appears at much higher levels in the apical membrane of the gut (Fig. 1F4). This gut-specific enrichment of the green signal was not observed with EGFP injections, suggesting that LvCMTM4 preferentially localizes to this region of the membrane or is stabilized there. In addition, CMTM4 protein appears to be apically localized in the gut (Fig. 1F4), suggesting that endogenous CMTM4 may be apically trafficked in this context.

To identify any functional roles for LvCMTM4 in sea urchin development, we performed a series of loss- and gain-of-function experiments (LOF and GOF, respectively). For LvCMTM4 LOF, we injected zygotes with a morpholino antisense oligonucleotide (MO) targeting LvCMTM4 and assessed the resulting embryos at 48 hours post fertilization (hpf). From morphological comparisons to control embryos at the same time point, LvCMTM4 MO-injected embryos are missing many skeletal elements, especially the secondary skeletal elements and ventral transverse rods (Fig. 2A-B, E). We also note that many MO-injected embryos appear stunted and/or have orientation defects in which the skeletal elements are abnormally rotated about the body axes (Fig. 2B, E). For LvCMTM4 GOF, we injected sea urchin zygotes with LvCMTM4 mRNA and scored the resulting embryos at 48 hpf. We found that LvCMTM4 mRNA-injected embryos are also missing many secondary skeletal elements and have dramatic rotational defects (Fig. 2C, F). Both perturbations produce a small fraction of embryos with spurious elements (Fig. 2E-F), although this is not a dominant phenotype. To confirm MO specificity, we co-injected LvCMTM4 MO with LvCMTM4 mRNA and scored the resulting embryos for normal or perturbed skeletal patterning (Fig. 2D, G). We found that while about half of MO- or mRNA-injected embryos have perturbed skeletal patterning, most of the co-injected embryos exhibit normal skeletal patterning. This rescue indicates that the CMTM4 MO is specific and lacks off-target effects.

**Figure 2.**
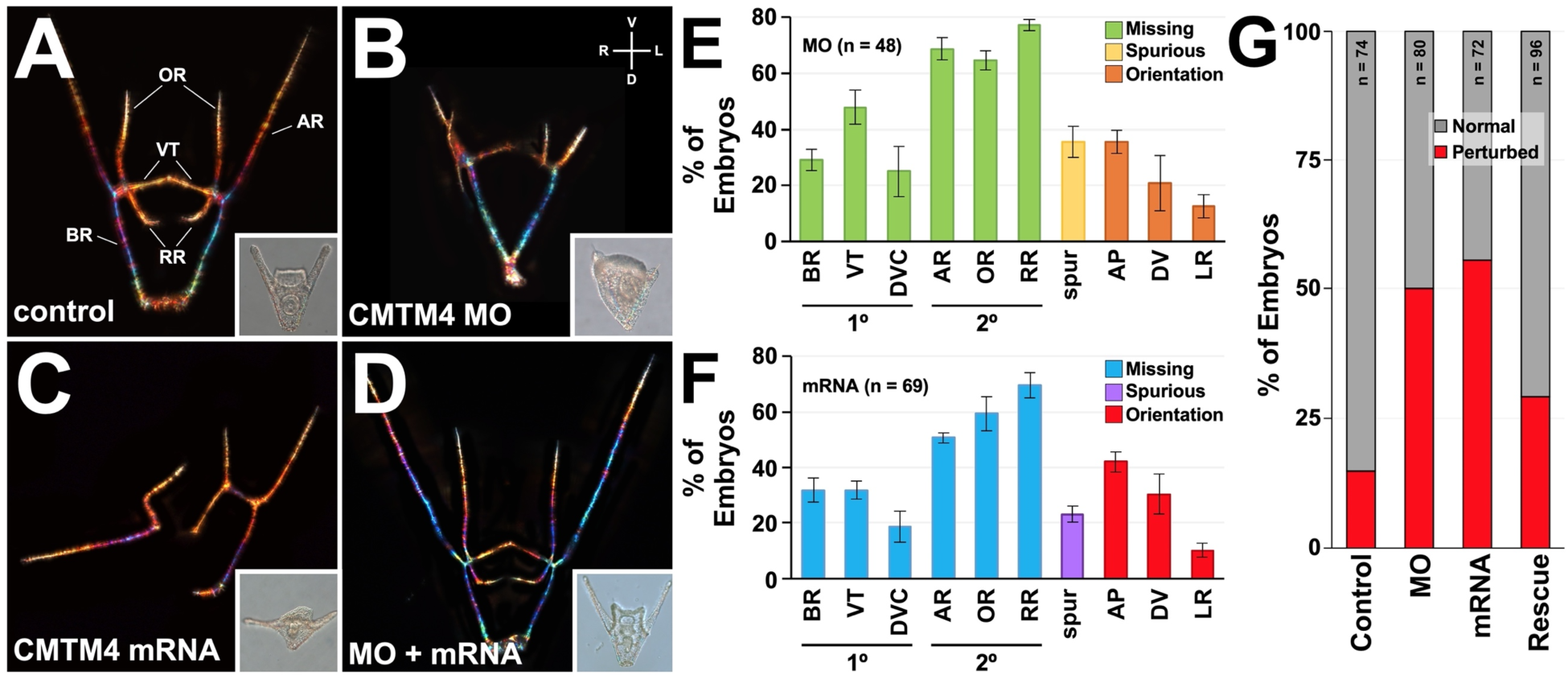
Normal levels of LvCMTM4 are required for normal skeletal patterning. **A-D.** Control (A), CMTM4 MO-injected (B), CMTM4 mRNA-injected (C), and CMTM4 MO and mRNA co-injected (D) embryos are shown at 48 hpf as skeletal birefringence images; insets show morphology (DIC) of the corresponding embryo. **E-F.** The percentages of CMTM4 MO-injected (E) and CMTM4 mRNA-injected (F) embryos with the indicated defects are shown as average ± S.E.M. The primary elements are the body rods (BR), ventral transverse rods (VTs), dorsal-ventral connecting rods (DVC); the secondary elements are the aboral rods (ARs), oral rods (ORs), and recurrent rods (RR). Other defects are spurious (spur) elements and orientation defects about the AP, DV, or LR body axes. **G.** The percentage of embryos with perturbed (red) or normal (grey) development in control, CMTM4 MO-injected (MO), CMTM4 mRNA-injected (mRNA), or CMTM4 MO and mRNA co-injected (Rescue) embryos at 48 hpf are shown as averages. Sample size (n) is indicated for each bar.

To test whether the CMTM4-EGFP fusion could phenocopy the CMTM4 GOF effects, we tested a range of CMTM4-EGFP mRNA doses. We found that all tested concentrations produced control-like embryos, including at 10-fold more concentrated than the effective dose of CMTM4 mRNA (Fig. S2). We confirmed that microinjection and gene expression were successful by the presence of C-terminal EGFP fluorescence (Fig. S2). Since the fusion construct appeared specifically localized *in vivo* and given that EGFP is a relatively large protein, this outcome suggests that fusion of EGFP to the C-terminus of CMTM4 sterically hinders its accessibility, implying that this intracellular terminus of CMTM4 is required for mediating its effects on skeletal patterning.

To assess the impact of CMTM4 perturbation on the spatiotemporal dynamics of biomineralization, we employed our polychrome labeling method for calcium detection (Descoteaux et al., 2023) using a simple two-fluorochrome labeling approach in which control, CMTM4 LOF, and CMTM4 GOF embryos were exposed to xylenol orange (XO) for the first 24 hours of development, then to calcein blue (CB) from 24 hpf until imaging at 48 hpf. 24 hpf was chosen as the switch time because the initial triradiate elements are typically completed by then in controls (Descoteaux et al., 2023). In the experiments herein, the triradiates, most of the body rods, and the bases of the aboral rods are well-labeled by XO prior to 24 hpf in controls (Fig. 3A). In comparison, CMTM4-perturbed embryos incorporated markedly less XO into their skeletons (Fig. 3B-C), indicating that biomineralization is delayed in both CMTM4 LOF and GOF embryos. It is unclear whether this reflects delayed initiation of the triradiates or slower progression of biomineralization after initiation. Interestingly, we also observe left-right asymmetries in the extent of XO labeling in CMTM4 LOF and GOF embryos (Fig. 3B-C). This suggests that biomineralization of the left and right skeletal spicules is desynchronized in CMTM4-perturbed embryos.

**Figure 3.**
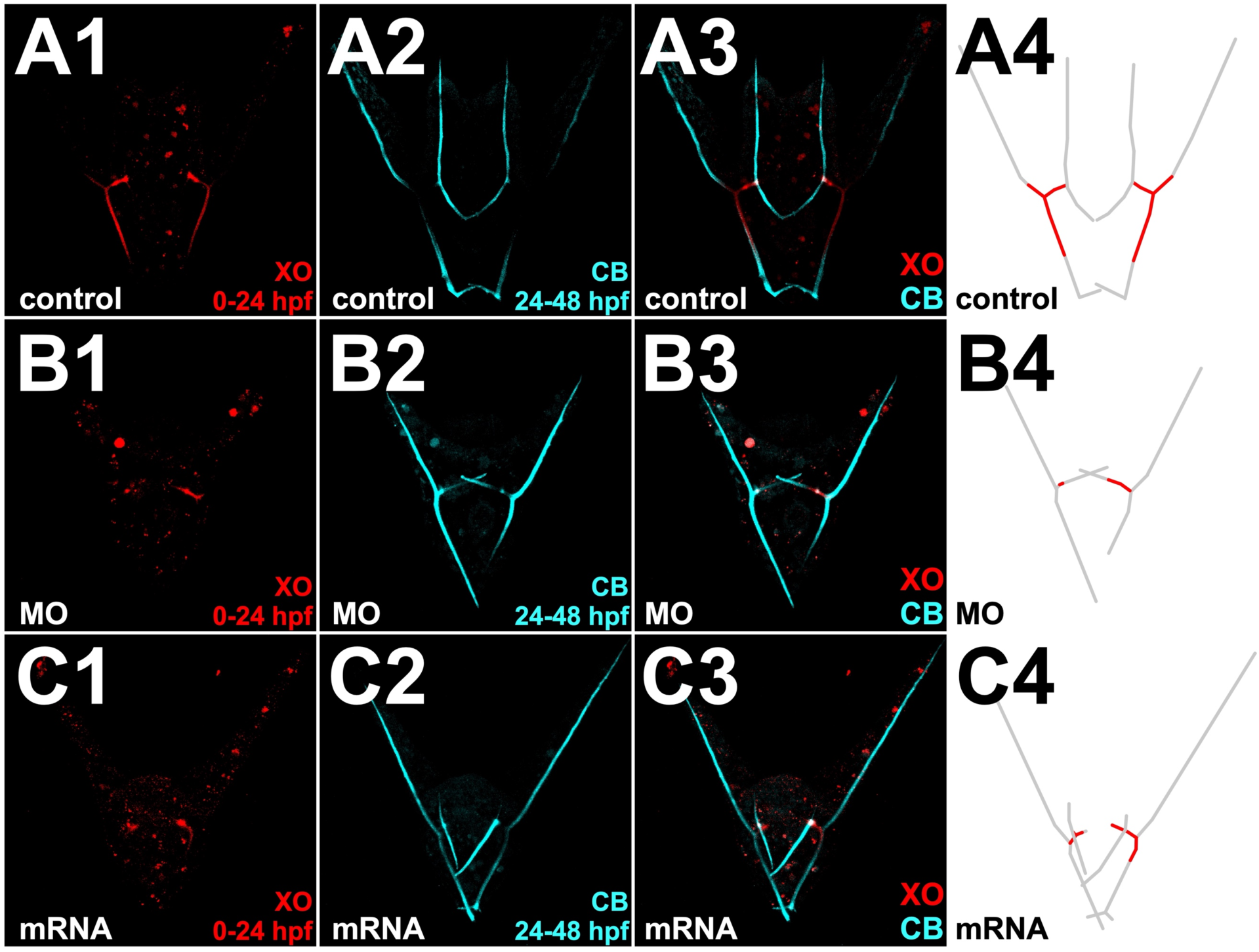
Polychrome labeling reveals temporal delay and left-right asynchrony in triradiate formation in LvCMTM4-perturbed embryos. Exemplar control (A), CMTM4 MO-injected (B), and CMTM4 mRNA-injected (C) embryo double-labeled as indicated are shown as individual fluorochromes (1-2) and merged (3). The extent of initial XO label incorporation (red) versus the entire skeleton (grey) is shown schematically (4). XO, xylenol orange; CB, calcein blue.

The skeletal biomineral is secreted by the PMCs into the lumen of their shared syncytial cable; therefore, the pattern of the larval skeleton is determined by the pattern of migration of the PMCs (Hodor and Ettensohn, 1998; Vidavsky et al., 2014; Wilt and Ettensohn, 2007; Wilt et al., 2008). Since CMTM4 affects cell migration in other systems (Bona et al., 2022; Chrifi et al., 2019; Xue et al., 2019), we were curious whether CMTM4 affects PMC migration in sea urchin larvae as well. To investigate this, we immunolabeled control, CMTM4 MO-injected, and CMTM4 mRNA-injected embryos at 48 hpf with a PMC-specific antibody (Fig. 4A-C) and scored for abnormalities in PMC migration and connectivity. We found that both CMTM4 LOF and GOF result in abnormal PMC migration, especially the failure of PMCs or clusters to migrate out of the ring-and-cords arrangement to pattern the secondary elements (Fig. 4B-D). Both perturbation conditions also produce ectopic PMCs, which are single PMCs that have migrated to abnormal locations and are disconnected from the PMC syncytium; this effect is significant with CMTM4 GOF, and more highly varied with LOF (Fig. 4D, Fig. S3). Some CMTM4 GOF embryos also have ectopic clusters of PMCs, though this defect is less prevalent in CMTM4 LOF embryos (Fig. 4D, Fig. S3). CMTM4 GOF also produces a significantly larger fraction of embryos with abnormal filopodial webbing (Fig. 4C, D). Breaks within the syncytial cable connecting PMCs within the normal pattern occur with CMTM4 LOF and, to a lesser extent, CMTM4 GOF (Fig. 4D). Taken together, these results show that normal levels of CMTM4 are required for normal PMC migration, syncytial integrity, and filopodial organization.

**Figure 4.**
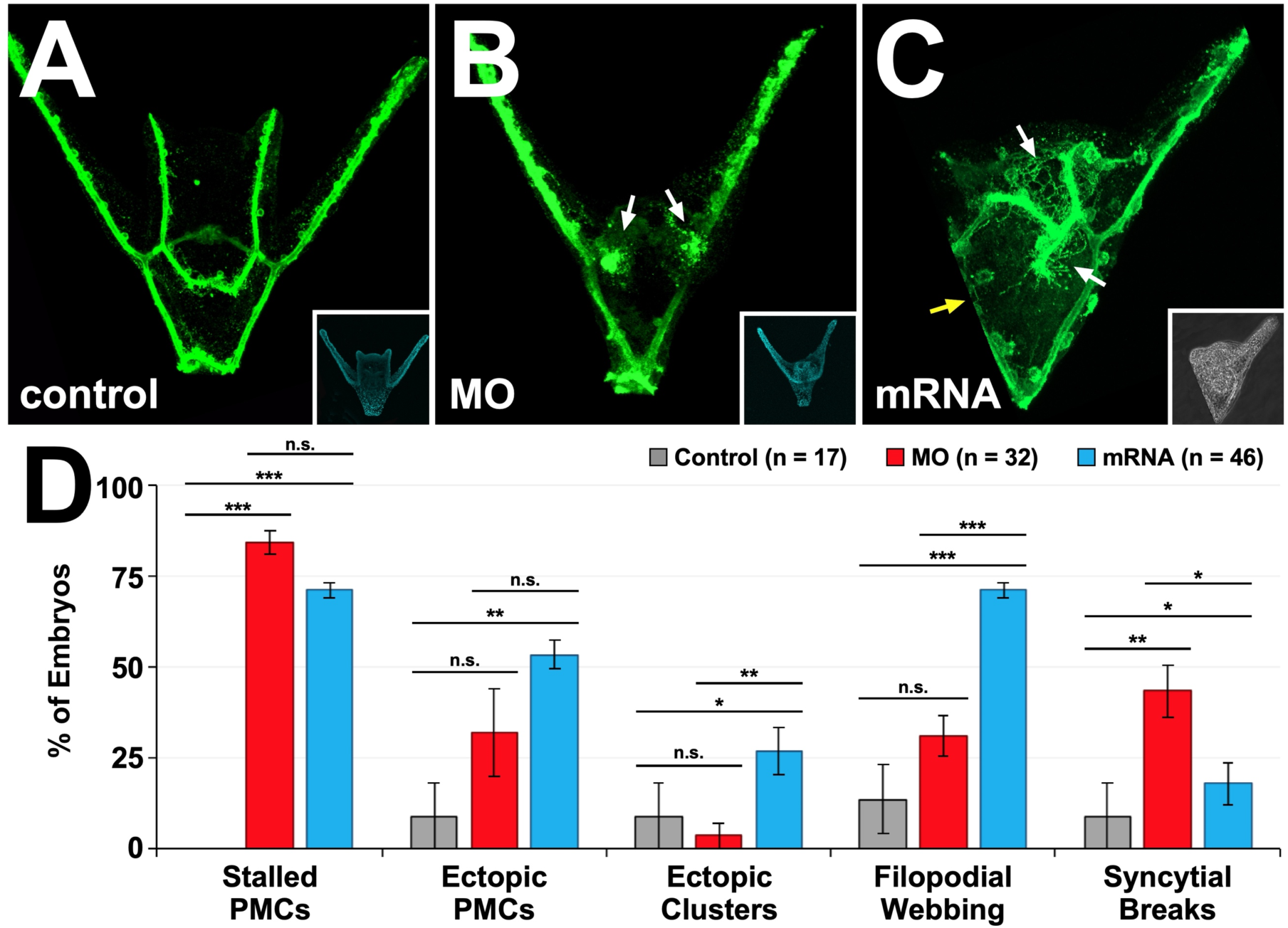
CMTM4 perturbation disrupts PMC migration and filopodial organization. **A-C.** Exemplar control (A), CMTM4 MO-injected (B), and CMTM4 mRNA-injected (C) 48 hpf embryos were immunolabeled with PMC-specific antibody 6a9. Insets show morphology via phase contrast (3) or nuclei labeled with Hoechst (1-2) in the corresponding embryo. Arrows in B indicate stalled PMC clusters; arrows in C indicate filopodial webbing (white) or syncytial breaks (yellow). **D.** The percentages of control (grey), CMTM4 MO-injected (red), or CMTM4 mRNA-injected (blue) embryos with the indicated PMC defects are shown as averages ± S.E.M. n.s. not significant; * p < 0.05; ** p < 0.005; *** p < 0.0005 (*t*-test).

The patterning of the primary mesenchyme cells (PMCs) is dictated by cues from the overlying ectoderm (Armstrong et al., 1993; Descoteaux et al., 2023; Ettensohn, 1990; Hardin and Armstrong, 1997; Hawkins et al., 2023; Malinda and Ettensohn, 1994; Piacentino et al., 2015; Piacentino et al., 2016a; Piacentino et al., 2016a; Piacentino et al., 2016b; Rodríguez-Sastre et al., 2023; Thomas et al., 2023; von Ubisch, 1937; Zuch and Bradham, 2019b). If the ectoderm is abnormally specified, the migration of the PMCs and subsequent skeletal pattern will be perturbed. A classic example is nickel treatment, which ventralizes the ectoderm and produces radialized skeletal patterns with supernumerary pentaradially-arranged triradiates (Hardin et al., 1992).

Because CMTM4-perturbed embryos exhibit abnormal skeletal patterning and PMC migration, we next investigated whether specification of the ectoderm is also affected. The ciliary band is a thin strip of ciliated cells that is spatially restricted to the ectodermal dorsal-ventral (DV) boundary by the DV specifying signals, making spatial restriction of the ciliary band an indicator for whether dorsal and ventral ectodermal specification have occurred normally (Bradham et al., 2009; Piacentino et al., 2016a; Yaguchi et al., 2010). We performed immunostains in control, CMTM4 LOF, and CMTM4 GOF embryos at 48 hpf to visualize the ciliated cells. We found that both CMTM4 LOF and CMTM4 GOF embryos have normal restriction of the ciliary region to a stripe approximately four cells wide (Fig. 5A-B). This suggests that ectodermal DV specification occurs normally in CMTM4-perturbed embryos.

**Figure 5.**
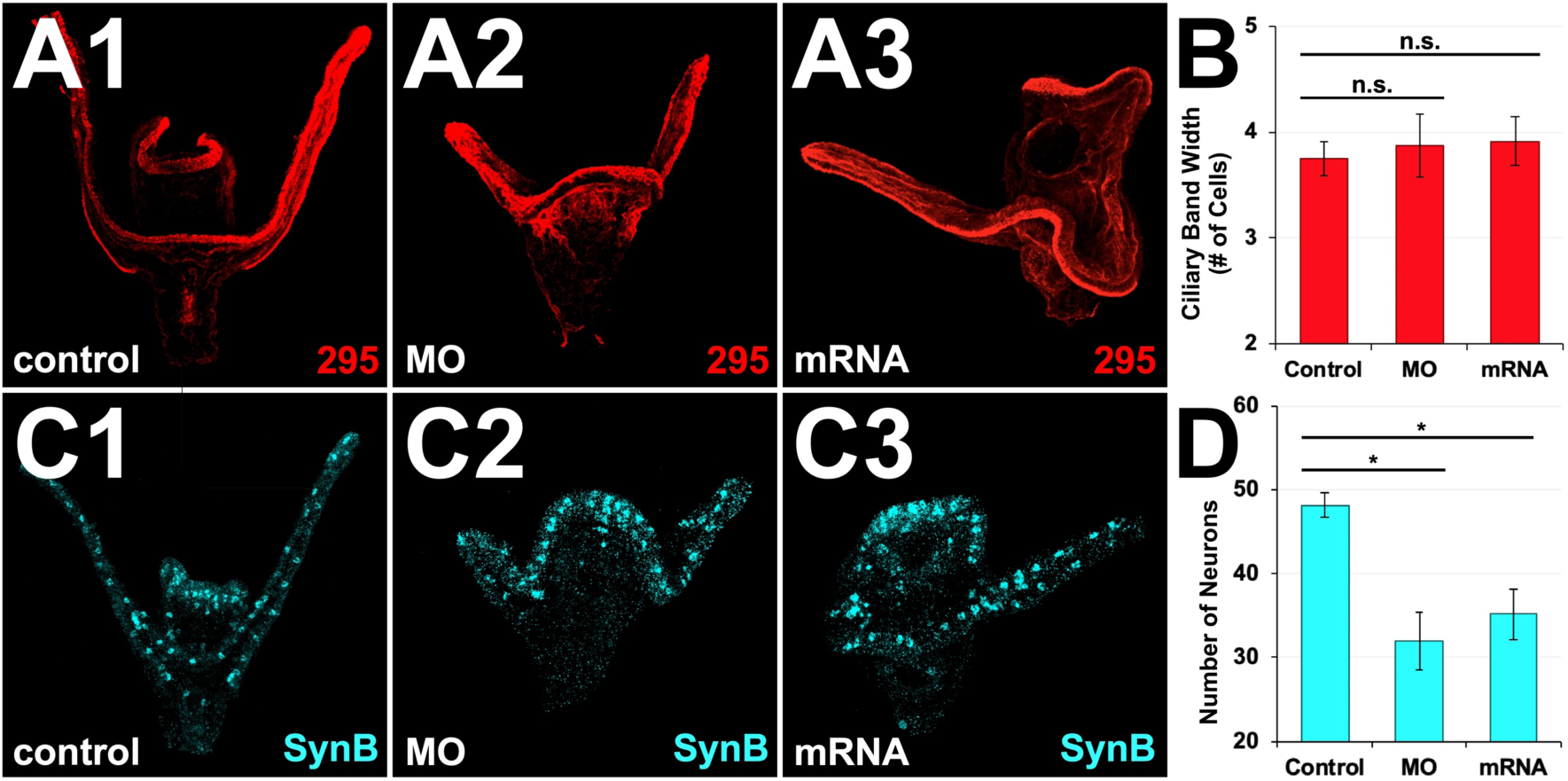
LvCMTM4 perturbation affects neuronal development but not overall dorsal-ventral ectodermal specification. **A.** Control (1), CMTM4 MO-injected (2), and CMTM4 mRNA-injected (3) embryos were fixed at 48 hpf and immunolabeled with ciliary band-specific antibody 295. **B.** The average width of the ciliary band is shown as average number of cells ± S.E.M. for each condition; n ≥ 8; n.s. not significant (*t*-test). **C.** Control (1), CMTM4 MO-injected (2), and CMTM4 mRNA-injected (3) 48 hpf embryos were subjected to HCR FISH for pan-neural gene synaptotagmin B (SynB). **D.** The average number of neurons in embryos from each condition is shown ± S.E.M. for each condition; n ≥ 6; * p < 0.05, *t*-test.

By pluteus stage, sea urchin larvae have also undergone neural development. The neurons are derived from the ectoderm and some innervate the ciliary band, directing swimming and feeding behaviors by regulating the beating of the cilia (Bradham et al., 2009; Mackie et al., 1969; Satterlie and Cameron, 1985; Strathmann, 2007; Yaguchi et al., 2010). Abnormalities in neural specification can also reflect disrupted ectodermal DV specification, making neuronal localization another readout for whether specification of these regions has occurred normally (Bradham et al., 2009; Piacentino et al., 2016a; Yaguchi et al., 2010). To visualize neural specification in CMTM4-perturbed embryos, we used HCR FISH for the pan-neural gene synaptotagmin B (SynB) in control, CMTM4 LOF, and CMTM4 GOF embryos at 48 hpf. We found that neurons are specified in similar locations in CMTM4 LOF and GOF embryos compared to controls (Fig. 5C); however, the number of neurons present in CMTM4 LOF or GOF embryos is significantly lower. On average, control embryos have approximately 48 neurons, while CMTM4 LOF and GOF embryos only have between 32 and 35 neurons, respectively (Fig. 5D). Because the neurons are in the correct location and the ciliary band is normally restricted, we conclude that CMTM4 perturbation is not sufficient to affect gross ectodermal DV specification. However, because the number of neurons specified in CMTM4-perturbed embryos is reduced compared to controls, we also conclude that normal CMTM4 levels are required for specification of a full complement of neurons.

Adhesion likely plays an important role in skeletal patterning. Throughout the patterning process, PMCs migrate both individually and collectively; PMCs in clusters exhibit larger numbers of contacts with other PMCs (≥ three contacts) compared to PMCs in the ring and cords, which form just two contacts. Finally, PMCs exhibit migratory behaviors that conclude with their positioning at specific spatial locations, implying adhesive interactions between the PMCs and the adjacent ectoderm. Thus, regulation of cell adhesion among the PMCs and between the PMCs and other cell types is probably an important aspect of skeletal patterning that is largely unexplored. The abnormally stalled and ectopic PMC clusters and syncytial breaks in CMTM4-perturbed embryos could therefore reflect abnormal regulation of cell-cell adhesion within the PMC clusters or between the PMCs and the ectoderm. To test whether LvCMTM4 influences PMC cell-cell adhesion, we devised a simple adhesion assay that takes advantage of the transcription factor Pmar1. Overexpression of *pmar1* via zygotic mRNA injection induces all cells to become PMCs (Oliveri et al., 2003). These PMCs dissociate from one another by 12 hpf and migrate radially in a random manner to produce a diffuse spread of similarly sized cells (Fig. 6A). However, the overexpression of an adhesion protein, such as Cadherin6 (Cad6), in pmar1-induced PMCs prevents these PMCs from dissociating and instead results in clumps of PMCs persisting even at 24 hpf (Fig. 6B). Interestingly, we found that *cmtm4* overexpression in pmar1-induced PMCs also resulted in a statistically significant fraction of PMC aggregates at 24 hpf, to a comparable extent as observed with *pmar1* and *cad6* co-injected embryos (Fig. 6C-D). Thus, we conclude that LvCMTM4 is sufficient to promote cell-cell adhesion of PMCs.

**Figure 6.**
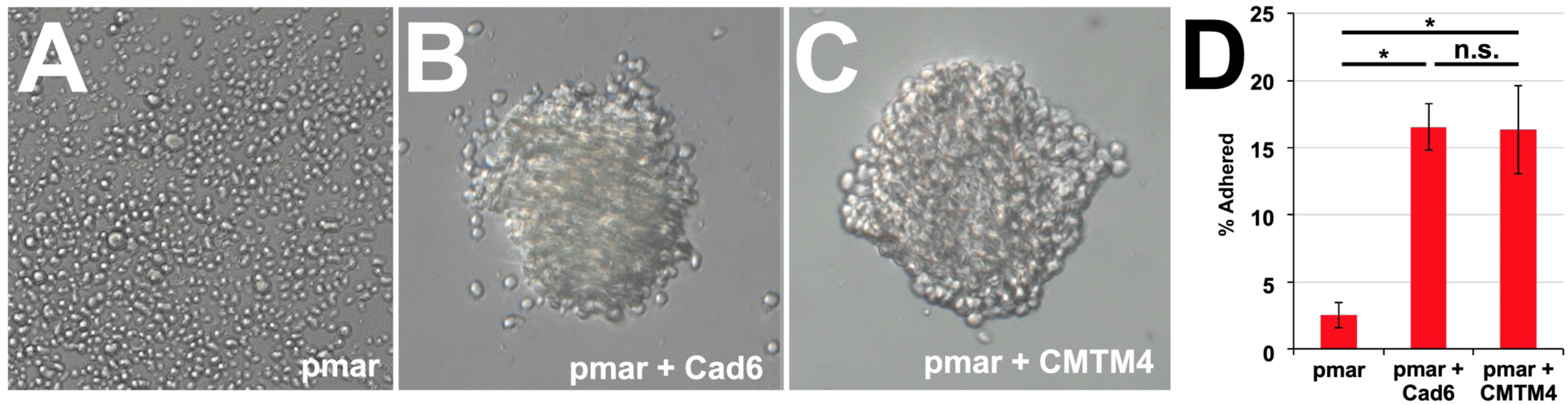
LvCMTM4 is sufficient to promote cell-cell adhesion in PMCs. **A-C.** Exemplar embryos injected with *pmar1* mRNA (A), *pmar1* and *cad6* mRNAs (B), and *pmar1* and *cmtm4* mRNAs (C). **D.** The percentages of pmar1-induced PMC clusters that remained adhered in clumps are shown as averages ± S.E.M. for *pmar1*-injected, *pmar1* and *cad6* co-injected, and *pmar1* and *cmtm4* co-injected embryos; n ≥ 16; * p < 0.05, n.s. not significant (*t*-test).

## Discussion

In this study, we characterize LvCMTM4 and its effects on skeletal patterning and biomineralization, PMC migration, and ectodermal specification. We find that LvCMTM4 is a typical MARVEL domain-containing protein with four transmembrane domains. The N- and C-termini are predicted to be intracellular, and our findings suggest that accessibility of the C-terminus is required for CMTM4 function. LvCMTM4 is globally expressed but appears to be elevated in the gut, particularly the apical membrane. The apical localization of CMTM4 is consistent with previous reports that MARVEL proteins concentrate at apical membranes (Puertollano and Alonso, 1999; Raleigh et al., 2010). The observed elevation of CMTM4 in the gut was unexpected for a potential regulator of skeletal patterning: our results imply that the gut may be a second source of skeletal patterning cues, along with the ectoderm. A role in the gut would also make sense if CMTM4 has other functions within the developing embryo. For example, the larval gut has important roles in uptake of food and other materials from the environment, such as via pinocytosis (Huvard and Holland, 1986; Strathmann, 1975). Because CMTM4 is known to regulate vesicular activity in other systems, it is possible that LvCMTM4 is playing a similar role here. The larval gut also plays a central role in coordinating the immune response to pathogens (Buckley et al., 2019; Ho et al., 2017). Previous studies have found that IL-17 is a critical component of the immune response in the gut of the sea urchin larva (Buckley et al., 2017). Since CMTM4 is known to regulate IL-17 signaling in other species (Knizkova et al., 2022), it is possible that LvCMTM4 is involved in the IL-17-mediated immune response in sea urchin larvae as well, which could explain the high level of LvCMTM4 expression in the larval gut. Another possibility is that LvCMTM4 plays a role in gut barrier function, which would make sense given the adhesive action of LvCMTM4 uncovered herein.

We report that CMTM4 knockdown and overexpression perturb skeletal patterning, resulting in the loss of many skeletal elements, particularly the secondary skeletal elements, as well as element orientation defects. This connection between CMTM4 and skeletal patterning is a novel observation. We note that the penetrance of defects in MO- and mRNA-injected embryos is somewhat low. MO injection is a knockdown approach, meaning that translation of the MO target will be inhibited in a dose-dependent manner. Given the lack of antibodies available to assess CMTM4 protein levels, we assume that the knockdown is partial and that some CMTM4 expression and activity persists. To confidently assess the effects of a complete loss of CMTM4 activity, a knockout approach will be needed. Recent advances have successfully developed genome editing protocols using the CRISPR/Cas9 system in sea urchins (Lin and Su, 2016; Nesbit et al., 2019; Wessel et al., 2020), which will facilitate knockout of genes involved in skeletal patterning such as CMTM4. As for the low penetrance of CMTM4 overexpression phenotypes, this suggests that compensatory mechanisms may be offsetting those effects. It is also likely that CMTM4 is serving multiple functions within the developing embryo and is not solely involved in skeletal patterning. Thus, although the overall skeletal pattern is unaffected by CMTM4 perturbation in some embryos, it is possible that other, less conspicuous processes not explored here, such as immune or gut barrier function, might be dysregulated by CMTM4 perturbation.

The fact that CMTM4 LOF and GOF produce the same rather than reciprocal defects is also interesting. This suggests that CMTM4 could be acting as a scaffold, such that multiple proteins dock on to CMTM4 to activate signaling cascades or other functions. A putative scaffolding function would likely involve one or both of its intracellular termini; that idea fits with our finding that sterically blocking the accessibility of the C-terminus of CMTM4 via fusion of EGFP prevents the CMTM4 overexpression phenotype. If CMTM4 functions as a scaffold protein, then too much CMTM4 as a result of overexpression would upset the scaffolding stoichiometry such that CMTM4’s binding partners are more likely to bind singly to CMTM4 copies rather than as pairs and thus fail to come together to interact, resulting in reduced CMTM4-dependent downstream activity. Similarly, too little CMTM4 as a result of CMTM4 knockdown would reduce or preclude scaffold formation by preventing CMTM4 binding proteins from interacting with each other and thereby block downstream processes. Additional work will be necessary to identify the putative binding partners that mediate CMTM4’s functions.

We also find that LvCMTM4 perturbation decreases and bilaterally desynchronizes biomineralization during the first 24 hours of development. The reduced biomineralization could reflect either a delay in initiation of skeletogenesis, as is observed in embryos perturbed by nickel treatment, or could reflect a slower rate of biomineralization, as is observed in VEGFR-inhibited embryos (Descoteaux et al., 2023). CMTM4 promotes endocytosis in other systems (Chrifi et al., 2019), and skeletal biomineralization relies on the uptake of calcium from the sea water via endocytosis (Hu et al., 2020; Mozingo, 2015; Vidavsky et al., 2014; Vidavsky et al., 2015). Dysregulation of CMTM4 activity could therefore result in reduced capacity for endocytosis, limiting the rate at which biomineralization that can occur. Further polychrome labeling experiments can use three-color approaches to measure the rate of elongation of the skeletal elements (Descoteaux et al., 2023), which could provide insight into whether the rate of biomineralization is indeed reduced in CMTM4-perturbed embryos.

CMTM4 perturbation disrupts PMC migration and filopodial organization. CMTM4 LOF and GOF both results in stalled PMC migration, in which clusters of PMCs fail to migrate to pattern the secondary skeletal elements. This indicates that LvCMTM4 is required for normal PMC migration in sea urchin larvae, which is consistent with previous findings that CMTM4 plays a role in regulating cell migration in other systems (Bona et al., 2022; Li et al., 2021; Xue et al., 2019). CMTM4 LOF also results in a high fraction of embryos with discontinuities in the PMC syncytium. This suggests that CMTM4 is required for syncytial integrity, perhaps via its adhesive function, which is a novel observation. In addition to stalled PMC migration, CMTM4 GOF also results in filopodial webbing, in which abnormal filopodial networks extend from both normally and ectopically located PMCs. The migration and patterning of the PMCs depends on instructive cues from the overlying ectoderm and the ability of the PMCs to correctly respond to those cues (Armstrong et al., 1993; Ettensohn, 1990; Hardin and Armstrong, 1997; Malinda and Ettensohn, 1994; Piacentino et al., 2015; Piacentino et al., 2016a; Piacentino et al., 2016b; Tan et al., 1998; Thomas et al., 2023; von Ubisch, 1937). The abnormal positioning of the PMCs and excessive filopodial networks in CMTM4 GOF embryos could therefore reflect a disruption of the ectodermal patterning cues that direct PMC migration and/or a loss of the ability of the PMCs to correctly respond to those patterning cues. Alternatively, these phenotypes could also reflect abnormal adhesion between the PMCs and ectoderm in CMTM4 perturbants. Although we did not assess whether CMTM4 mediates cell-cell adhesion between PMCs and ectoderm, this function is plausible given the broad expression of CMTM4.

Our results also show that the PMC and skeletal patterning defects observed following CMTM4 LOF or GOF are likely not due to gross defects in dorsal-ventral ectodermal specification, since the ciliary band is normally restricted and the neurons are correctly positioned in CMTM4-perturbed embryos. This makes sense, since the skeletal abnormalities observed following CMTM4 perturbation are distinct from the patterning defects observed in embryos with a mis-specified ectoderm, such as radialized skeletons (Hardin et al., 1992). However, this does not rule out the possibility that the ectoderm is more subtly affected by CMTM4 perturbation, resulting in the reduction in neuronal numbers observed with CMTM4 morphants. It is possible that CMTM4 perturbation results in changes in ectodermal gene expression, such as the loss of patterning cues, that would contribute to the ectopic and/or stalled migration of the PMCs and subsequent abnormal skeletal patterning observed in CMTM4-perturbed embryos. Analysis of the changes in gene expression in CMTM4 LOF and GOF embryos compared to controls, such as through RNA sequencing, will be necessary to clarify this.

We also observed that although CMTM4 perturbation does not affect the location of neurons, it does reduce the number of neurons that develop in the larvae. Whether this reflects differential specification in CMTM4 perturbants, or differential neuronal survival therein remains unclear. The reduced number of neurons could possibly be due to the abnormal morphologies of CMTM4-perturbed embryos, rather than issues with neuronal specification itself. CMTM4-perturbed embryos are often missing one or more of the four arms normally present in the 48 hpf pluteus-stage larval skeleton. Since these arms are typically innervated, the absence of one or more arms could reduce the total number of neurons present in the embryo, since neurons are not needed to innervate that area. Perhaps such neurons are unable to find targets and thus do not survive.

Finally, we find that CMTM4 overexpression promotes cell-cell adhesion of pmar1-induced PMCs. These findings are consistent with previous reports in other systems that CMTM4 plays a role in regulating the cohesiveness of cell clusters during cell migration (Bona et al., 2022; Chrifi et al., 2019); however, whether CMTM4 is directly involved in the anchoring of the cells to one another is unclear. Apical expression of CMTM4 is consistent with it mediating a direct adhesive contact. However, in other systems, CMTM4 regulates the level of cell surface expression of adhesion proteins such as Cadherin (Chrifi et al., 2019). It is therefore possible that the increased rate of PMC-PMC adhesion observed following CMTM4 overexpression is a result of increased localization of Cadherin to the cell surface rather than direct participation of CMTM4 in cell adhesion complexes. Further work will be needed to make this distinction, and to determine whether CMTM4 also mediates adhesion between PMCs and other cell types. Nonetheless, our work is the first to identify a gene, CMTM4, as a regulator of cohesiveness between PMCs during sea urchin skeletal patterning. Overall, this work establishes LvCMTM4 as a novel regulator of PMC migration and adhesion, skeletal patterning, and neuronal development, and sets the stage for additional studies to uncover the mechanisms by which CMTM4 provokes its effects.

## Materials and Methods

### Animals and embryo cultures

Adult *Lytechinus variegatus* sea urchins were obtained from the Duke University Marine Laboratory (Beaufort, NC), Laura Salter (Davis, NC), or Reeftopia (Miami, FL). Gamete harvesting, fertilization, and embryo culturing was performed as previously described (Bradham and McClay, 2006). At least two biological replicates were collected for each experiment.

### Microinjections, morpholinos, and constructs

Microinjections were performed as previously described (Bradham and McClay, 2006; Piacentino et al., 2016a). PCR products containing the Lv-CMTM4 and Lv-Pmar1 open reading frames were cloned into pCS2+ for mRNA synthesis. pCS2-CMTM4-EGFP and pCS2-EGFP constructs were synthesized by GenScript (Piscataway, NJ), while pCS2-LvCadherin6 construct was synthesized by Twist Bioscience (San Francisco, CA). The pCS2-memb-mCherry construct was obtained from Addgene (Addgene #53750)(Megason, 2009). All injected mRNAs were transcribed *in vitro* using the mMessage mMachine kit (Ambion). The translation-blocking CMTM4 MO was obtained from GeneTools. The MO sequence is 5’-GCAAAGTGATGTTTGCAGTGTCAGACATCTTTAAATCT-3’. Dose-response experiments were performed with all reagents to determine their optimal working concentrations. Unless otherwise specified, injection concentrations were 0.27-0.4 mM for CMTM4 MO, 400 ng/μL (phenotyping) or 1500 ng/μL (adhesion assay) for CMTM4 mRNA, 100 ng/μL for *pmar1* mRNA, 2000 ng/μL for *cadherin6* mRNA, and 50 ng/μL for CMTM4-EGFP, EGFP, and memb-mCherry mRNA.

### Skeletal imaging and scoring

Embryos were imaged on a Zeiss Axioplan microscope at 200x magnification with differential interference contrast (DIC) to visualize morphology or at multiple focal planes with plane-polarized light to capture skeletal birefringence. Montaged skeletal images were produced with ImageJ and excess out-of-focus light was manually removed to view the entire skeleton in focus. All focal planes were used for skeletal scoring using an in-house scoring rubric (Piacentino et al., 2016a; Piacentino et al., 2016b; Thomas et al., 2023). For phenotypic counts, embryos were photographed in large groups with DIC imaging at 50x magnification, then scored and counted from single focus images (Thomas et al., 2023).

### Polychrome labeling

Xylenol orange tetrasodium salt (Sigma #398187) and calcein blue (Sigma #M1255) stocks were prepared by dissolving each chemical in distilled water. Embryos were incubated in 30 μM xylenol orange (XO) or 45 μM calcein blue (CB). Two-color fluorochrome labeling experiments were performed as described in (Descoteaux et al., 2023), with color switches occurring at 24 hpf.

### Immunolabels, HCR FISH, and confocal microscopy

Embryos were fixed in 4% paraformaldehyde in artificial sea water (ASW) at 48 hpf prior to immunolabeling or HCR FISH. Immunolabeling was performed as previously described (Bradham et al., 2009). Ciliary band labeling was performed with undiluted monoclonal 295 primary antibody (a gift from David McClay, Duke University, Durham, NC). PMC labeling was performed with monoclonal 6a9 primary antibody (1:30; a gift from Charles Ettensohn, Carnegie Mellon University, Pittsburgh, PA). Hoechst 33258 (Sigma #94403) was used at 1:1000 to label nuclei. Fluorescent secondary antibodies goat anti-mouse Alexa 488 (Invitrogen #A11001) or goat anti-mouse DyLight™ 405 (Jackson Laboratories #115-475-003) were used at 1:500 dilution.

Hybridization chain reaction fluorescent *in situ* hybridization (HCR FISH) probe sets were designed from the open reading frames for LvCMTM4, LvVEGFR, and LvSynB by Molecular Instruments, Inc. (Los Angeles, CA). Probe sets, buffers, and amplifier hairpins fluorescently labeled with Alexa 488, Alexa 546, or Alexa 647 were obtained from Molecular Instruments, Inc. Embryos were fixed at 48 hpf and subjected to HCR FISH protocol for sea urchin embryos (Choi et al., 2018; Choi et al., 2020). All steps were performed in 1.5 mL microcentrifuge tubes. Embryos were incubated in hairpin solutions for at least 3h in the dark at room temperature, then washed with 5X SSCT before mounting in 50% PBS/glycerol for imaging.

Confocal imaging was performed using an Olympus Fv10i laser-scanning confocal microscope or a Nikon C2Si laser-scanning confocal microscope. Z-stacks were collected at 400x and used to generate maximum-intensity projections using ImageJ. Cell counts, such as ciliary band width and total neurons, were obtained manually from maximum-intensity projections.

### Adhesion assay

*L. variegatus* zygotes were microinjected with *pmar1* mRNA alone as a negative control or were co-injected either with *pmar1* and *cadherin6* mRNAs or with *pmar1* and *cmtm4* mRNAs. Injected zygotes were immediately transferred into 35 mm glass-bottom imaging dishes with 14 mm micro-wells containing ASW (CellVis # D35-14-1.5-N). Embryos were imaged at 24 hpf and the fraction of PMC clusters that remained adhered was calculated and averaged across two biological replicates each for pmar alone and pmar with Cad6 co-injections and three biological replicates for the pmar and CMTM4 co-injections.

## Acknowledgements

We would like to thank Todd Blute for equipment training and support. We would also like to thank Professors Charles Ettensohn and David McClay for antibodies and Dan Zuch for the LvPmar1 construct. M.R. would like to thank the Boston University Trustee Scholarship Program.

## Author contributions

Conceptualization: C.A.B.; Methodology: A.E.D., C.A.B.; Investigation: A.E.D., M.R., D.A.; Formal analysis: A.E.D., M.R., C.A.B.; Writing-original draft: A.E.D; Writing – review & editing: A.E.D., C.A.B.; Funding acquisition: C.A.B.

## References

Armstrong, N., Hardin, J. and McClay, D. R. (1993). Cell-cell interactions regulate skeleton formation in the sea urchin embryo. Development 119, 833–840.

Bona, A., Seifert, M., Thünauer, R., Zodel, K., Frew, I. J., Römer, W., Walz, G. and Yakulov, T. A. (2022). MARVEL domain containing CMTM4 affects CXCR4 trafficking. Mol. Biol. Cell 33, ar116.

Bradham, C. A. and McClay, D. R. (2006). p38 MAPK is essential for secondary axis specification and patterning in sea urchin embryos. Development 133, 21–32.

Bradham, C. A., Oikonomou, C., Kühn, A., Core, A. B., Modell, J. W., McClay, D. R. and Poustka, A. J. (2009). Chordin is required for neural but not axial development in sea urchin embryos. Dev. Biol. 328, 221–233.

Buckley, K. M., Ho, E. C. H., Hibino, T., Schrankel, C. S., Schuh, N. W., Wang, G. and Rast, J. P. (2017). IL17 factors are early regulators in the gut epithelium during inflammatory response to Vibrio in the sea urchin larva. eLife 6, e23481.

Buckley, K. M., Schuh, N. W., Heyland, A. and Rast, J. P. (2019). Analysis of immune response in the sea urchin larva. In Methods in Cell Biology, pp. 333–355. Elsevier.

Choi, H. M. T., Schwarzkopf, M., Fornace, M. E., Acharya, A., Artavanis, G., Stegmaier, J., Cunha, A. and Pierce, N. A. (2018). Third-generation *in situ* hybridization chain reaction: multiplexed, quantitative, sensitive, versatile, robust. Development 145, dev165753.

Choi, H. M. T., Schwarzkopf, M. and Pierce, N. A. (2020). Multiplexed Quantitative In Situ Hybridization with Subcellular or Single-Molecule Resolution Within Whole-Mount Vertebrate Embryos: qHCR and dHCR Imaging (v3.0). In In Situ Hybridization Protocols (ed. Nielsen, B. S.) and Jones, J.), pp. 159–178. New York, NY: Springer US.

Chrifi, I., Louzao-Martinez, L., Brandt, M. M., van Dijk, C. G. M., Bürgisser, P. E., Changbin Zhu, Changbin Zhu, Zhu, C., Zhu, C., Kros, J. M., et al. (2019). CMTM4 regulates angiogenesis by promoting cell surface recycling of VE-cadherin to endothelial adherens junctions. Angiogenesis 22, 75–93.

Descoteaux, A. E., Zuch, D. T. and Bradham, C. A. (2023). Polychrome labeling reveals skeletal triradiate and elongation dynamics and abnormalities in patterning cue-perturbed embryos. Dev. Biol. 498, 1–13.

Ettensohn, C. A. (1990). The regulation of primary mesenchyme cell patterning. Dev. Biol. 140, 261–271.

Ettensohn, C. A. (2017). Sea urchins as a model system for studying embryonic development. In Reference Module in Biomedical Sciences, p. B9780128012383995096. Elsevier.

Furuse, M., Hirase, T., Itoh, M., Nagafuchi, A., Yonemura, S., Tsukita, S. and Tsukita, S. (1993). Occludin: a novel integral membrane protein localizing at tight junctions. J. Cell Biol. 123, 1777–1788.

Gustafson, T. and Wolpert, L. (1963). The cellular basis of morphogenesis and sea urchin development. Int. Rev. Cytol.-Surv. Cell Biol. 15, 139–214.

Gustafson, T. and Wolpert, L. (1967). Cellular movement and contact in sea urchin morphogenesis. Biol. Rev. 42, 442–498.

Haass, N. K., Kartenbeck, M. A. and Leube, R. E. (1996). Pantophysin is a ubiquitously expressed synaptophysin homologue and defines constitutive transport vesicles. J. Cell Biol. 134, 731–746.

Hallgren, J., Tsirigos, K. D., Pedersen, M. D., Almagro Armenteros, J. J., Marcatili, P., Nielsen, H., Krogh, A. and Winther, O. (2022). DeepTMHMM predicts alpha and beta transmembrane proteins using deep neural networks.

Han, W., Ding, P., Xu, M., Wang, L., Rui, M., Shi, S., Liu, Y., Zheng, Y., Chen, Y., Yang, T., et al. (2003). Identification of eight genes encoding chemokine-like factor superfamily members 1–8 (CKLFSF1–8) by in silico cloning and experimental validation. Genomics 81, 609–617.

Hardin, J. and Armstrong, N. (1997). Short-Range Cell–Cell Signals Control Ectodermal Patterning in the Oral Region of the Sea Urchin Embryo. Dev. Biol. 182, 134–149.

Hardin, J., Coffman, J. A., Black, S. D. and McClay, D. R. (1992). Commitment along the dorsoventral axis of the sea urchin embryo is altered in response to NiCl2. Development 116, 671–685.

Hawkins, D. Y., Zuch, D. T., Huth, J., Rodriguez-Sastre, N., McCutcheon, K. R., Glick, A., Lion, A. T., Thomas, C. F., Descoteaux, A. E., Johnson, W. E., et al. (2023). ICAT: a novel algorithm to robustly identify cell states following perturbations in single-cell transcriptomes. Bioinformatics 39, btad278.

Ho, E. C., Buckley, K. M., Schrankel, C. S., Schuh, N. W., Hibino, T., Solek, C. M., Bae, K., Wang, G. and Rast, J. P. (2017). Perturbation of gut bacteria induces a coordinated cellular immune response in the purple sea urchin larva. Immunol. Cell Biol. 95, 647–647.

Hodor, P. G. and Ettensohn, C. A. (1998). The dynamics and regulation of mesenchymal cell fusion in the sea urchin embryo. Dev. Biol. 199, 111–124.

Hu, M. Y., Petersen, I., Chang, W. W., Blurton, C. and Stumpp, M. (2020). Cellular bicarbonate accumulation and vesicular proton transport promote calcification in the sea urchin larva. Proc. R. Soc. B Biol. Sci. 287, 20201506.

Huvard, A. L. and Holland, N. D. (1986). Pinocytosis of ferritin from the gut lumen in larvae of a sea star (*Patiria miniata*) and a sea urchin (*Lytechinus pictus*). Dev. Growth Differ. 28, 43–51.

Ikenouchi, J., Furuse, M., Furuse, K., Sasaki, H., Tsukita, S. and Tsukita, S. (2005). Tricellulin constitutes a novel barrier at tricellular contacts of epithelial cells. J. Cell Biol. 171, 939–945.

Jin, C., Ding, P., Wang, Y. and Ma, D. (2005). Regulation of EGF receptor signaling by the MARVEL domain-containing protein CKLFSF8. FEBS Lett. 579, 6375–6382.

Jumper, J., Evans, R., Pritzel, A., Green, T., Figurnov, M., Ronneberger, O., Tunyasuvunakool, K., Bates, R., Žídek, A., Potapenko, A., et al. (2021). Highly accurate protein structure prediction with AlphaFold. Nature 596, 583–589.

Knizkova, D., Pribikova, M., Draberova, H., Semberova, T., Trivic, T., Synackova, A., Ujevic, A., Stefanovic, J., Drobek, A., Huranova, M., et al. (2022). CMTM4 is a subunit of the IL-17 receptor and mediates autoimmune pathology. Nat. Immunol. 23, 1644–1652.

Li, H., Li, J., Su, Y., Fan, Y., Guo, X., Li, L., Su, X., Rong, R., Ying, J., Mo, X., et al. (2014). A novel 3p22.3 gene CMTM7 represses oncogenic EGFR signaling and inhibits cancer cell growth. Oncogene 33, 3109–3118.

Li, H., Liu, Y.-T., Chen, L., Zhou, J.-J., Chen, D.-R., Li, S.-J. and Sun, Z.-J. (2021). CMTM4 regulates epithelial-mesenchymal transition and PD-L1 expression in head and neck squamous cell carcinoma. Mol. Carcinog. 60, 556–566.

Lin, C.-Y. and Su, Y.-H. (2016). Genome editing in sea urchin embryos by using a CRISPR/Cas9 system. Dev. Biol. 409, 420–428.

Mackie, G. O., Spencer, A. N. and Strathmann, R. (1969). Electrical activity associated with ciliary reversal in an echinoderm larva. Nature 223, 1384–1385.

Malinda, K. M. and Ettensohn, C. A. (1994). Primary mesenchyme cell migration in the sea urchin embryo: Distribution of directional cues. Dev. Biol. 164, 562–578.

Malinda, K. M., Fisher, G. W. and Ettensohn, C. A. (1995). Four-dimensional microscopic analysis of the filopodial behavior of primary mesenchyme cells during gastrulation in the sea urchin embryo. Dev. Biol. 172, 552–566.

Megason, S. G. (2009). In Toto Imaging of Embryogenesis with Confocal Time-Lapse Microscopy. In Zebrafish (ed. Lieschke, G. J.), Oates, A. C.), and Kawakami, K.), pp. 317–332. Totowa, NJ: Humana Press.

Mezzadra, R., Sun, C., Jae, L. T., Gomez-Eerland, R., de Vries, E., Wu, W., Wu, W., Logtenberg, M. E. W., Slagter, M., Rozeman, E. A., et al. (2017). Identification of CMTM6 and CMTM4 as PD-L1 protein regulators. Nature 549, 106–110.

Miller, J. R., Fraser, S. E. and McClay, D. R. (1995). Dynamics of thin filopodia during sea urchin gastrulation. Development 121, 2501–2511.

Mozingo, N. M. (2015). Lectin uptake and incorporation into the calcitic spicule of sea urchin embryos. Zygote 23, 467–473.

Nesbit, K. T., Fleming, T., Batzel, G., Pouv, A., Rosenblatt, H. D., Pace, D. A., Hamdoun, A. and Lyons, D. C. (2019). The painted sea urchin, Lytechinus pictus, as a genetically-enabled developmental model. In Methods in Cell Biology, pp. 105–123. Elsevier.

Oliveri, P., Davidson, E. H. and McClay, D. R. (2003). Activation of pmar1 controls specification of micromeres in the sea urchin embryo. Dev. Biol. 258, 32–43.

Peterson, R. E. and McClay, D. R. (2003). Primary mesenchyme cell patterning during the early stages following ingression. Dev. Biol. 254, 68–78.

Piacentino, M. L., Ramachandran, J. and Bradham, C. A. (2015). Late Alk4/5/7 signaling is required for anterior skeletal patterning in sea urchin embryos. Development 142, 943–952.

Piacentino, M. L., Zuch, D. T., Fishman, J., Rose, S., Sviatlana Rose, Speranza, E., Li, C., Yu, J., Chung, O., Ramachandran, J., et al. (2016a). RNA-Seq identifies SPGs as a ventral skeletal patterning cue in sea urchins. Development 143, 703–714.

Piacentino, M. L., Chung, O., Ramachandran, J., Zuch, D. T., Yu, J., Conaway, E. A., Reyna, A. and Bradham, C. A. (2016b). Zygotic LvBMP5-8 is required for skeletal patterning and for left-right but not dorsal-ventral specification in the sea urchin embryo. Dev. Biol. 412, 44–56.

Plate, M., Li, T., Ting Li, Li, T., Yu Wang, Wang, Y., Mo, X., Zhang, Y., Ma, D. and Han, W. (2010). Identification and characterization of CMTM4, a novel gene with inhibitory effects on HeLa cell growth through Inducing G2/M phase accumulation. Mol. Cells 29, 355–361.

Puertollano, R. and Alonso, M. A. (1999). MAL, an integral element of the apical sorting machinery, is an itinerant protein that cycles between the *trans* -Golgi network and the plasma membrane. Mol. Biol. Cell 10, 3435–3447.

Raleigh, D. R., Marchiando, A. M., Zhang, Y., Shen, L., Sasaki, H., Wang, Y., Long, M. and Turner, J. R. (2010). Tight junction–associated MARVEL proteins MarvelD3, tricellulin, and occludin have distinct but overlapping functions. Mol. Biol. Cell 21, 1200–1213.

Rodríguez-Sastre, N., Shapiro, N., Hawkins, D. Y., Lion, A. T., Peyreau, M., Correa, A. E., Dionne, K. and Bradham, C. A. (2023). Ethanol exposure perturbs sea urchin development and disrupts developmental timing. Dev. Biol. 493, 89–102.

Sánchez-Pulido, L., Martín-Belmonte, F., Valencia, A. and Alonso, M. A. (2002). MARVEL: a conserved domain involved in membrane apposition events. Trends Biochem. Sci. 27, 599–601.

Satterlie, R. A. and Cameron, A. R. (1985). Electrical activity at metamorphosis in larvae of the sea urchin *Lytechinus pictus*. J. Exp. Zool. 235, 197–204.

Schatzberg, D., Lawton, M. L., Hadyniak, S. E., Ross, E. J., Carney, T., Beane, W. S., Levin, M. and Bradham, C. A. (2015). H(+)/K(+) ATPase activity is required for biomineralization in sea urchin embryos. Dev. Biol. 406, 259–270.

Strathmann, R. R. (1975). Larval feeding in echinoderms. Am. Zool. 15, 717–730.

Strathmann, R. R. (2007). Time and extent of ciliary response to particles in a non-filtering feeding mechanism. Biol. Bull. 212, 93–103.

Stumpp, M., Hu, M. Y., Melzner, F., Gutowska, M. A., Dorey, N., Himmerkus, N., Holtmann, W. C., Dupont, S., Thorndyke, M. C. and Bleich, M. (2012). Acidified seawater impacts sea urchin larvae pH regulatory systems relevant for calcification. Proc. Natl. Acad. Sci. U. S. A. 109, 18192–18197.

Takeuchi, H., Konnai, S., Maekawa, N., Minato, E., Ichikawa, Y., Kobayashi, A., Okagawa, T., Murata, S. and Ohashi, K. (2020). Expression Analysis of Canine CMTM6 and CMTM4 as Potential Regulators of the PD-L1 Protein in Canine Cancers. Front. Vet. Sci. 7, 330.

Tan, H., Ransick, A., Wu, H., Dobias, S. L., Liu, Y.-H. and Maxson, R. E. (1998). Disruption of primary mesenchyme cell patterning by misregulated ectodermal expression of SpMsx in sea urchin embryos. Dev. Biol. 201, 230–246.

The UniProt Consortium, Bateman, A., Martin, M.-J., Orchard, S., Magrane, M., Ahmad, S., Alpi, E., Bowler-Barnett, E. H., Britto, R., Bye-A-Jee, H., et al. (2023). UniProt: the Universal Protein Knowledgebase in 2023. Nucleic Acids Res. 51, D523–D531.

Thomas, C. F., Hawkins, D. Y., Skidanova, V., Marrujo, S. R., Gibson, J., Ye, Z. and Bradham, C. A. (2023). Voltage-gated sodium channel activity mediates sea urchin larval skeletal patterning through spatial regulation of Wnt5 expression. Development 150, dev201460.

Vidavsky, N., Addadi, S., Mahamid, J., Shimoni, E., Ben-Ezra, D., Shpigel, M., Weiner, S. and Addadi, L. (2014). Initial stages of calcium uptake and mineral deposition in sea urchin embryos. Proc. Natl. Acad. Sci. U. S. A. 111, 39–44.

Vidavsky, N., Masic, A., Schertel, A., Weiner, S. and Addadi, L. (2015). Mineral-bearing vesicle transport in sea urchin embryos. J. Struct. Biol. 192, 358–365.

von Ubisch (1937). Die normale Skelettbildung bei Echinodyamus pusillus and Psammechinus miliaris. Z. Für Wiss. Zool. 149, 402–476.

Wessel, G. M., Kiyomoto, M., Shen, T.-L. and Yajima, M. (2020). Genetic manipulation of the pigment pathway in a sea urchin reveals distinct lineage commitment prior to metamorphosis in the bilateral to radial body plan transition. Sci. Rep. 10, 1973.

Wilt, F. H. and Ettensohn, C. A. (2007). The Morphogenesis and Biomineralization of the Sea Urchin Larval Skeleton. In Handbook of Biomineralization (ed. Bäuerlein, E.), pp. 182–210. Wiley.

Wilt, F. H., Killian, C. E., Hamilton, P. and Croker, L. (2008). The dynamics of secretion during sea urchin embryonic skeleton formation. Exp. Cell Res. 314, 1744–1752.

Wolpert, L. and Gustafson, T. (1961). Studies on the cellular basis of morphogenesis of the sea urchin embryo. Development of the skeletal pattern. Exp. Cell Res. 25, 311–325.

Wu, S.-Y., Ferkowicz, M. J., Michael J. Ferkowicz and McClay, D. R. (2007). Ingression of primary mesenchyme cells of the sea urchin embryo: a precisely timed epithelial mesenchymal transition. Birth Defects Res. Part C Embryo Today Rev. 81, 241–252.

Xue, H., Ting Li, Li, T., Wang, P., Mo, X., Zhang, H., Ding, S., Ma, D., Lv, W., Jing Zhang, et al. (2019). CMTM4 inhibits cell proliferation and migration via AKT, ERK1/2, and STAT3 pathway in colorectal cancer. Acta Biochim. Biophys. Sin. 51, 915–924.

Yaguchi, S., Yaguchi, J., Angerer, R. C., Angerer, L. M. and Burke, R. D. (2010). TGFβ signaling positions the ciliary band and patterns neurons in the sea urchin embryo. Dev. Biol. 347, 71–81.

Yuan, W., Liu, B., Wang, X., Li, T., Xue, H., Mo, X., Yang, S., Ding, S. and Han, W. (2017). CMTM3 decreases EGFR expression and EGF-mediated tumorigenicity by promoting Rab5 activity in gastric cancer. Cancer Lett. 386, 77–86.

Yuan, Y., Sheng, Z., Liu, Z., Zhang, X., Xiao, Y., Xie, J., Zhang, Y. and Xu, T. (2020). CMTM5-v1 inhibits cell proliferation and migration by downregulating oncogenic EGFR signaling in prostate cancer cells. J. Cancer 11, 3762–3770.

Zacchetti, D., Peränen, J., Murata, M., Fiedler, K. and Simons, K. (1995). VIP17/MAL, a proteolipid in apical transport vesicles. FEBS Lett. 377, 465–469.

Zuch, D. T. and Bradham, C. A. (2019a). Spatially mapping gene expression in sea urchin primary mesenchyme cells. Methods Cell Biol. 151, 433–442.

Zuch, D. T. and Bradham, C. A. (2019b). Spatially mapping gene expression in sea urchin primary mesenchyme cells. Methods Cell Biol. 151, 433–442.

