## Supplemental Figures for "CMTM4 is an adhesion modulator that regulates skeletal patterning and primary mesenchyme cell migration in sea urchin embryos"

**Descoteaux et al.**

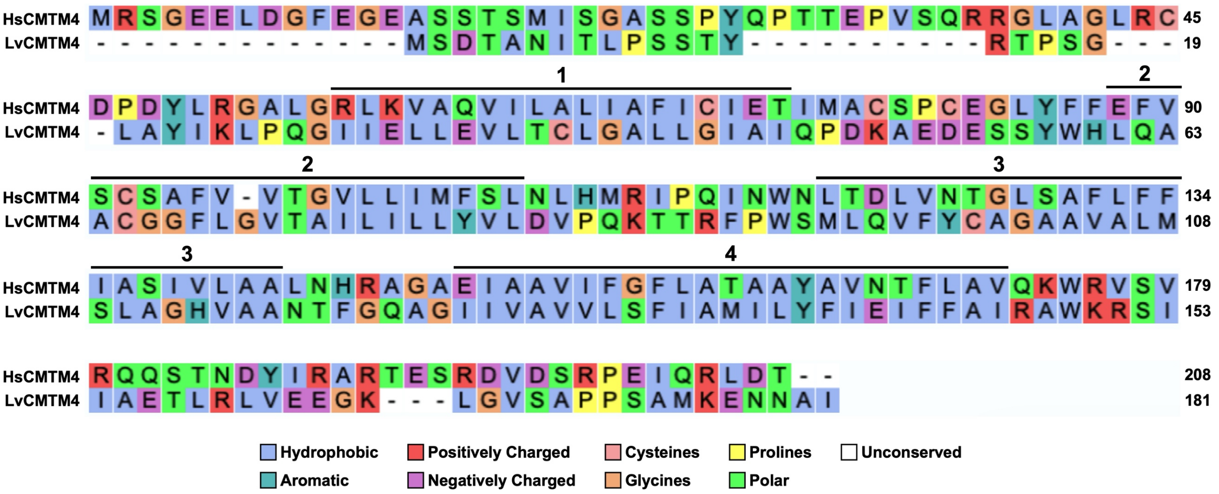


**Figure S1. Clustal alignment of *H. sapiens* and *L. variegatus* CMTM4 protein sequences.** Residues are color-coded according to the Clustal2 default color scheme. The four predicted LvCMTM4 transmembrane domains are indicated by black lines above the peptide sequence.

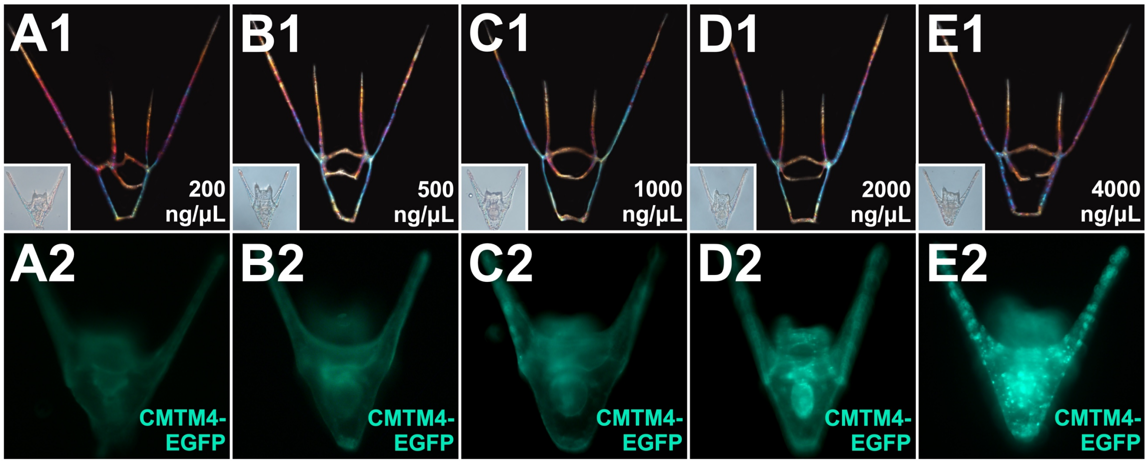


**Figure S2. LvCMTM4-EGFP overexpression is not sufficient to induce skeletal patterning defects. A-E.** Exemplar embryos injected with 200 (A), 500 (B), 1000 (C), 2000 (D), or 4000 ng/µL (E) of LvCMTM4-EGFP mRNA are shown at 48 hpf as skeletal birefringence (1) and epifluorescence (2) images. Insets shows morphology (DIC) of the corresponding embryos. Compare to Fig. 2C.

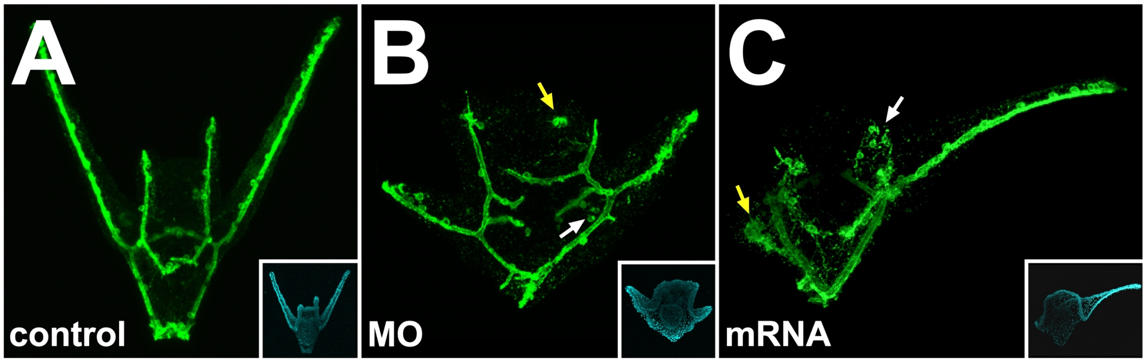


**Figure S3. CMTM4 perturbation results in ectopic PMCs and clusters. A-C.** Exemplar control (A), CMTM4 MO-injected (B), and CMTM4 mRNA-injected (C) 48 hpf embryos were immunolabeled with PMC-specific antibody 6a9. Insets show nuclei labeled with Hoechst in the corresponding embryo. Arrows in B-C indicate ectopic PMCs (white) or ectopic clusters (yellow).
